# Early-life hypoxia alters adult physiology and reduces stress resistance and lifespan in *Drosophila*

**DOI:** 10.1101/2020.03.31.018960

**Authors:** Danielle M Polan, Mohammad Alansari, Byoungchun Lee, Savraj Grewal

**Author notes:** These authors contributed equally (Danielle M Polan, Mohammad Alansari, Byoungchun Lee).

## Abstract

In many animals, short-term fluctuations in environmental conditions in early life often exert long-term effects on adult physiology. In Drosophila, one ecologically relevant environmental variable is hypoxia. *Drosophila* larvae live on rotting, fermenting food rich in microorganisms – an environment characterized by low ambient oxygen. They have therefore evolved to tolerate hypoxia. While the acute effects of hypoxia in larvae have been well studied, whether early-life hypoxia affects adult physiology and fitness is less clear. Here we show that Drosophila exposed to hypoxia during their larval period subsequently show reduced starvation stress resistance and shorter lifespan as adults, with these effects being stronger in males. We find that these effects are associated with reduced whole-body insulin signaling but elevated TOR kinase activity, a manipulation known to reduce lifespan. We also identify a sexually dimorphic effect of larval hypoxia on adult nutrient storage and mobilization. Thus, we find that males, but not females, showing elevated levels of lipids and glycogen. Moreover, we see that both males and females exposed to hypoxia as larvae show defective lipid mobilization upon starvation stress as adults. These data show how early-life hypoxia can exert persistent, sexually dimorphic, long-term effects on Drosophila adult physiology and lifespan.

## INTRODUCTION

Animals often live in conditions where environmental conditions such as temperature, food, oxygen, and pathogen exposure, can fluctuate dramatically. The ability of animals to adapt their metabolism and physiology to these changing environments is essential for their survival. Many adaptive responses occur immediately in response to changes in environment, particularly in response to environmental stressors (e.g. starvation, hypoxia, infection), to allow animals to survive whilst subjected to these stress conditions. It is also increasingly appreciated that acute, early-life environmental stresses can trigger longer-term responses that can influence later adult physiology and fitness (Gluckman and Hanson, 2004; Burdge and Lillycrop, 2014). In some cases, these early life environmental changes can confer subsequent beneficial effects on adult fitness. For example, starvation stress in larval honeybees leads to subsequent starvation tolerance as adults (Wang et al., 2016b; Wang et al., 2016a). In a similar manner, anoxia exposure during the development of the Caribbean fruit fly confers later anoxia resistance in adults (Visser et al., 2018). Early life mild heat stress in the zebra finch has also been shown to lower oxidative damage induced by heat stress in adult life (Costantini et al., 2012). In contrast to these adaptive responses, in some situations early life environmental stress can have deleterious consequences on subsequent adult physiology. Examples of these types of responses have been described in rodents where prenatal exposure to a deficient maternal diet subsequently leads to cardiovascular and metabolic defects, and shortened lifespan in adults (Langley-Evans et al., 1999; Aihie Sayer et al., 2001; Woods et al., 2001). These effects are examples of a concept known as the developmental origins of health and disease (DOHaD), which proposes that poor intra-uterine conditions during fetal development (often caused by defective maternal nutrition) can subsequently increase risk of metabolic disease in adulthood (Bruce and Hanson, 2010; Hanson and Gluckman, 2014). This hypothesis is supported by many epidemiological studies in humans showing that low birth weight (a proxy for poor intra-uterine environment) is associated with a number of metabolic diseases such as diabetes, obesity and heart disease (Gluckman et al., 2008). Together, these various reports emphasize the importance of investigating the mechanisms by which different early life environmental stresses can alter adult physiology.

Drosophila has been an excellent model to study how environmental cues influence physiology, development and lifespan. In particular, several recent reports have described how modulation in environment during the larval period of the life cycle can subsequently influence adult physiology and aging. For example, when Drosophila larvae are raised on low nutrients they subsequently show an extension of adult lifespan (Stefana et al., 2017). These effects were mediated by secretion of lipid autotoxic pheromones in adults. In other studies, when Drosophila were subjected to mild oxidative stress only during the larval period, this led to microbiome remodelling and persistent epigenetic changes that led to an extension of adult lifespan (Borch Jensen et al., 2017; Obata et al., 2018). Together, these studies how altered early larval life environmental conditions can cause persistent and long-lasting effects of on adult Drosophila physiology.

An important environmental variable in the Drosophila life cycle is oxygen exposure. In their natural ecology, Drosophila larvae grow by burrowing into rotting, fermenting food that is rich in microorganisms (Markow, 2015). This environment is likely low in oxygen and, as a result, Drosophila have evolved mechanisms to tolerate hypoxia. For example, when exposed to moderate (5-10% oxygen) hypoxia in the laboratory, larvae slow their growth and development, but can maintain their viability (Harrison and Haddad, 2011; Heinrich et al., 2011; Callier et al., 2015; Lee et al., 2019). These adaptive effects are mediated through several different changes in larval physiology including increased tracheal branching, changes in cell-cell signaling and metabolic gene expression, and altered lipid metabolism (Wingrove and O’Farrell, 1999; Centanin et al., 2008; Zhou et al., 2008; Li et al., 2013; Zhou and Haddad, 2013; Wong et al., 2014; Lee et al., 2019). These changes have been shown to allow larvae to survive and maintain homeostasis whilst exposed to low oxygen. However, whether larval hypoxia exposure exerts any persistent, long-term effects on adult physiology is not entirely clear. We explore this question in this paper.

## MATERIALS AND METHODS

### *Drosophila* stocks

All experiments were performed using *w*^*1118*^ flies. Flies were kept on medium containing 150 g agar, 1600 g cornmeal, 770 g Torula yeast, 675 g sucrose, 2340 g D-glucose, 240 ml acid mixture (propionic acid/phosphoric acid) per 34 L water and maintained at 25°C.

### Hypoxia exposure

For all hypoxia experiments *Drosophila* larvae were exposed to 5% oxygen. This was achieved by placing vials containing *Drosophila* into an airtight glass chamber into which a mix of 5% oxygen/95% nitrogen continually flowed. Flow rate was controlled using an Aalborg model P gas flow meter.

### Measurement of *Drosophila* starvation stress and lifespan

Eggs were collected on grape plates for 3-4 hours and then the next day, hatched larvae were transferred to vials (50 larvae per vial). Newly hatched larvae were then placed into one of two experimental conditions (see Figure 1): NORMOXIC experimental condition – larvae maintained in normoxia until adulthood; HYPOXIC experimental condition - larvae were maintained in hypoxia chambers for the duration of their larval period. They were then transferred back to normoxia as pupae allowed to develop to adulthood. For both experimental conditions, eclosed flies were allowed to mate for two days, and then males and females were separated under light anaesthesia into cohorts of 20 flies per vial. For the starvation stress experiments, flies were transferred at 5-6 post-eclosion into vials containing only 0.8% agar/PBS. Viability was then assessed twice daily until all flies had died. For the lifespan experiments, flies were transferred into fresh vials every 2-3 days and the number of dead flies counted until all flies had died.

**Fig 1.**
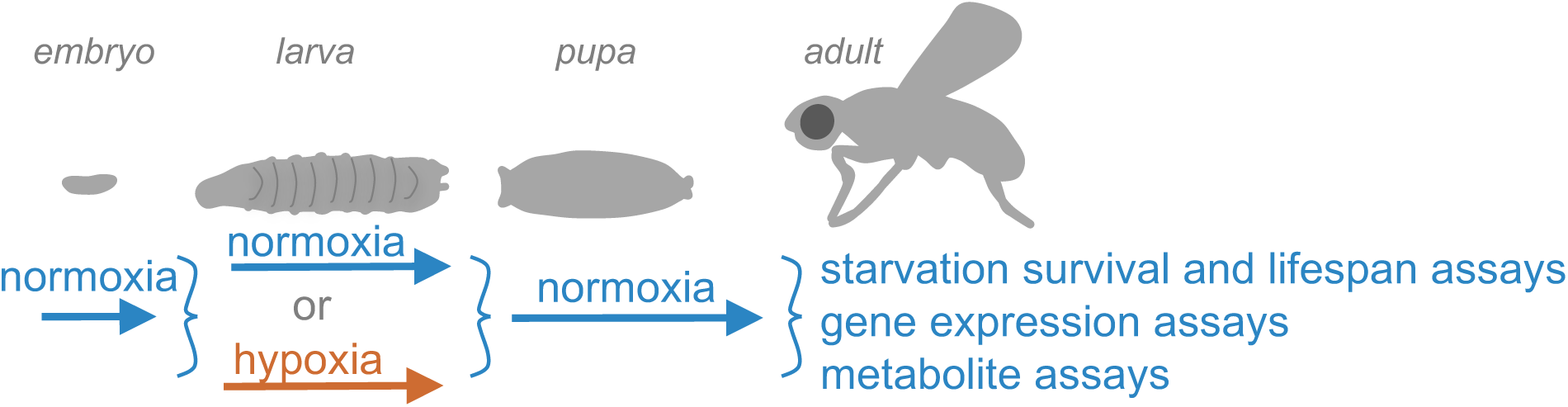
An outline of the experimental protocol. An outline of the experimental plan for examining the effects of larval hypoxia on adult physiology. For all experiments, *w*^*1118*^ embryos were raised in normoxia. Upon hatching, they were transferred to food vials and then kept in either normoxia (referred to in this paper as the NORMOXIC condition) or hypoxia (5% oxygen - referred to as the HYPOXIC condition) for the duration of their larval period. The animals were then kept in normoxia throughout pupal development until they emerged as adults. Mated, one-week old adults were then assayed for changes in their starvation stress survival, lifespan, gene expression, metabolite levels.

### qPCR analyses

Total RNA was extracted from groups of 5 adults using TRIzol reagent according to manufacturer’s instructions (Invitrogen; 15596–018). The RNA samples were treated with DNase (Ambion; 2238 G) and then reverse transcribed using Superscript II (Invitrogen; 100004925). The cDNAs were then used as a template for subsequent qRT–PCRs using SyBr Green PCR mix and an ABI 7500 real time PCR system. The PCR data were normalized to actin mRNA levels. The following primers were used:

*Actin5C* forward: GAGCGCGGTTACTCTTTCAC

*Actin5C* reverse: GCCATCTCCTGCTCAAAGTC

*dILP2* forward: TCCACAGTGAAGTTGGCCC

*dILP2* reverse: AGATAATCGCGTCGACCAGG

*dILP3* forward: AGAGAACTTTGGACCCCGTGAA

*dILP3* reverse: TGAACCGAACTATCACTCAACAGTCT

*dILP5* forward: GAGGCACCTTGGGCCTATTC

*dILP5* reverse: CATGTGGTGAGATTCGGAGCTA

### Western blotting

Groups of five adult *Drosophila* were lysed in a buffer containing 20 mM Tris-HCl (pH 8.0), 137 mM NaCl, 1 mM EDTA, 25 % glycerol, 1% NP-40, 50 mM NaF, 1 mM PMSF, 1 mM DTT, 5 mM sodium ortho vanadate (Na_3_VO_4_) and Protease Inhibitor cocktail (Roche Cat. No. 04693124001) and Phosphatase inhibitor (Roche Cat. No. 04906845001). Protein concentrations were measured using the Bio-Rad Dc Protein Assay kit II (5000112). Protein lysates (15 μg to 30μg) were resolved by SDS–PAGE and electro transferred to a nitrocellulose membrane, and then subjected to Western blotting with specific primary antibodies and HRP-conjugated secondary antibodies, and then visualized by chemiluminescence (enhanced ECL solution (Perkin Elmer). Primary antibodies used in this study were: anti-phospho-S6K-Thr398 (1:1000, Cell Signalling Technology #9209), anti-pAkt-T342 (gift from Michelle Bland, 1:1000 dilution), anti-pAkt-S505 (Cell Signaling #4054, 1:1000 dilution) and anti-actin (1:1000, Santa Cruz Biotechnology, # sc-8432). Secondary antibodies were purchased from Santa Cruz Biotechnology (sc-2030, 2005, 2020, 1:10,000 dilution).

### Metabolite measurements

Groups of five flies were weighed and then frozen in Eppendorf tubes on dry ice. Total glycogen and TAG levels were determined using colorimetric assays following the protocols described in detail in (Tennessen et al., 2014). For TAG assays, animals were lysed and lysates were heated at 70 Celsius for 10 minutes. Then they were incubated first with triglyceride reagent (Sigma; T2449) and then mixed with free glycerol reagent (Sigma; F6428). Colorimetric measurements were then made using absorbance at 540 nm and TAG levels calculated by comparing with a glycerol standard curve. Glycogen assays were performed by lysing animals in PBS and then heating lysates at 70 Celsius for 10 minutes. For each experimental sample, duplicate samples were either treated with amyloglucosidase (Sigma A1602) to breakdown glycogen intro glucose, or left untreated, and then levels of glucose in both duplicates measured by colorimetric assay following the addition of a glucose oxidase reagent (Sigma; GAGO-20). Levels of glycogen in each experimental sample were then calculated by subtracting the glucose measurements of the untreated duplicate from the amyloglucosidase-treated sample. All experimental metabolite concentrations were calculated by comparison with glycogen and glucose standard curves. All calculated metabolite levels were then corrected for adult body weight.

### Statistical analyses

Lifespan and stress survival data were analyzed using a Long-rank test. Metabolite and qPCR data were analyzed by Students t-test. All statistical analyses and data plots were performed using Prism software.

## RESULTS

The outline for all experiments is shown in Figure 1A. We chose to examine the effects of 5% oxygen exposure because at this level of oxygen larvae show reduced growth and altered metabolism, but still maintain normal food intake and develop into viable adults (Lee et al., 2019). For all experiments, *w*^*1118*^ embryos were raised in normoxia and when they hatched they were transferred to food vials and then maintained in either normoxia or hypoxia for the duration of their larval period. The animals were then kept in normoxia throughout pupal development until they emerged as adults. Mated, one-week old adults were then assayed for changes in their physiology caused by prior larval hypoxia (referred to in this paper as the HYPOXIC condition) compared to adults raised in normoxia (referred to as the NORMOXIC condition).

### Larval hypoxia leads to reduced adult starvation stress tolerance and shorter lifespan

We first examined whether larval exposure to hypoxia could subsequently alter adult stress responses and lifespan. We first compared the ability of the HYPOXIC and NORMOXIC adults to tolerate starvation stress. We found that compared to NORMOXIC animals, the HYPOXIC condition adults showed a significant reduction in viability when completely deprived of nutrients (Figure 2A, B). This reduced starvation tolerance was more pronounced in male (29.4% decrease in median survival in HYPOXIC flies) compared to females (5.6% decrease in median survival in HYPOXIC flies). We next examined the effects of larval hypoxia on adult lifespan. We found that HYPOXIC animals had a reduced lifespan compared to the NORMOXIC animals (Figure 2C, D). As with the starvation responses, these reductions in lifespan in HYPOXIC animals were stronger in males (17.6% decrease in median lifespan) compared to females (6.9% decrease in median lifespan). Together these results indicate that when exposed to hypoxia as larva, adult Drosophila have a reduced lifespan and a reduced ability to tolerate starvation.

**Fig 2.**
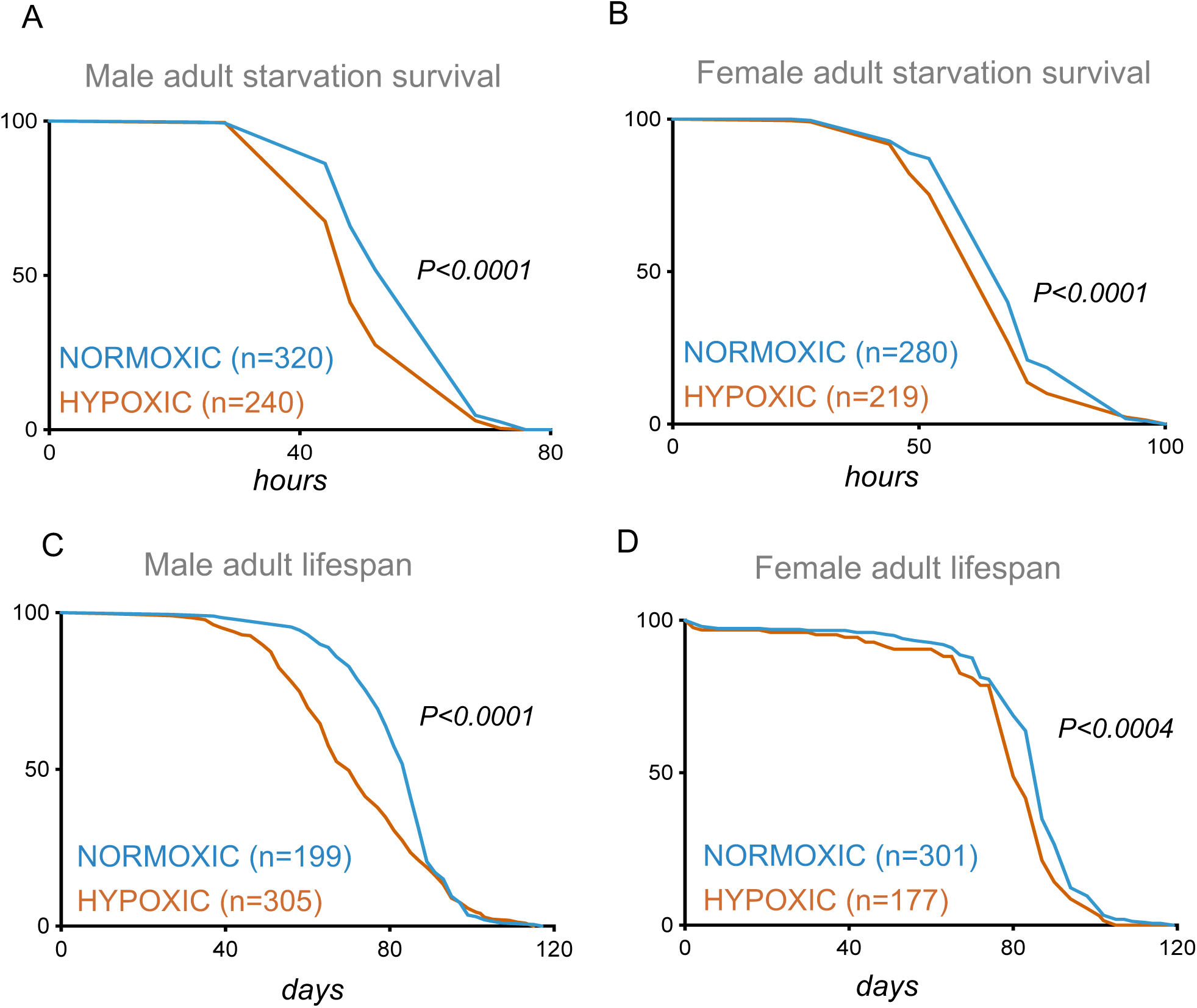
Larval hypoxia leads to reduced adult tolerance to starvation stress and reduced adult lifespan. (A, B) Data represent starvation survival curves for (A) male and (B) female adult Drosophila that had been exposed to either normoxia (blue lines) or hypoxia (orange lines) as larvae. Data were analyzed using the Log-rank test. (C, D) Data represent survival curves for (C) male and (D) female adult Drosophila that had been exposed to either normoxia (blue lines) or hypoxia (orange lines) as larvae. Data were analyzed using the Log-rank test.

### Larval hypoxia leads to decreased adult insulin signaling but increased TOR signaling

The conserved insulin and TOR kinase signalling pathways are major regulators of systemic metabolism and physiology in Drosophila. In particular, both signaling pathways have been shown to control stress responses in adults. For example, genetic or pharmacological lowering of either insulin or TOR signalling has been shown to extend lifespan and to increase tolerance to different stresses including starvation and oxidative stress (Katewa and Kapahi, 2011; Partridge et al., 2011). Given the effects of larval hypoxia on adult lifespan and stress tolerance that we observed, we examined whether larval exposure to hypoxia could lead to altered insulin or TOR signaling in adults.

We first measured mRNA levels of Drosophila insulin-like peptides (dILPs). Drosophila contain seven main dILPs that can bind the insulin receptor and activate a conserved downstream PI3K/Akt kinase pathway (Nassel et al., 2015). In particular, three dILPs (2, 3 and 5) that are expressed and secreted from neurosecretory cells in the larval and adult brains have been shown to influence stress responses and lifespan in Drosophila (Nassel and Vanden Broeck, 2016). When we measured dILP levels by qPCR, we found that the HYPOXIC adults showed reduced expression of dILP 2, 3 and 5 compared to the NORMOXIC animals (Figure 3A, B). We then measured phosphorylation of Akt in whole body lysates as a read-out for systemic insulin signalling. When the insulin pathway is activated, Akt is phosphorylated at two sites, threonine 342 and serine 505. We found that phosphorylation at both these sites was reduced in the HYPOXIC adults compared to NORMOXIC controls (Figure 3C).

**Fig 3.**
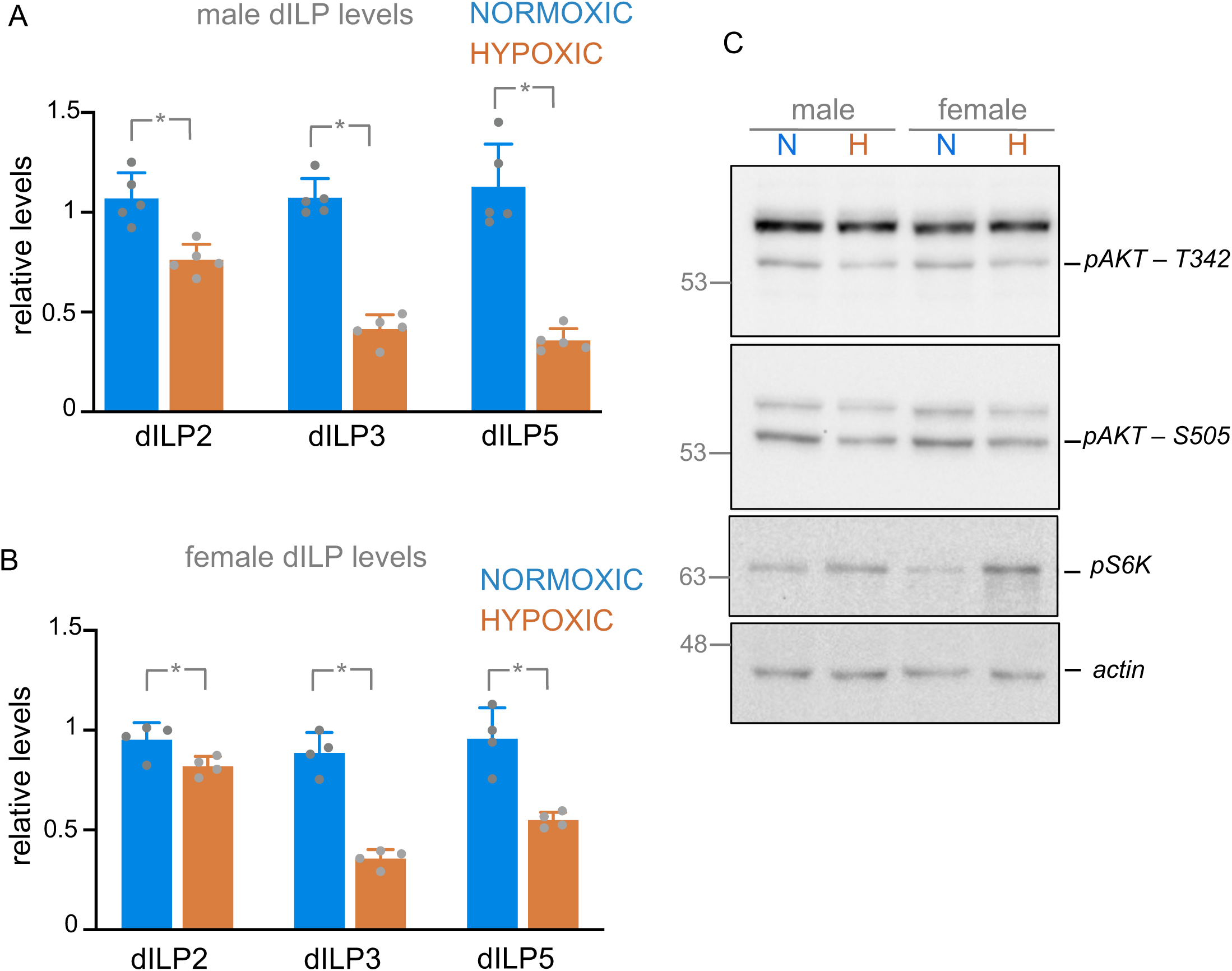
Larval hypoxia leads altered adult insulin and TOR signaling. A, B) Levels of dILP mRNAs were measured by qRT-PCR in mated adult males (A) and females (B) after they had been exposed to either normoxia (blue plots) or hypoxia (orange plots) as larvae. Data are presented as mean + standard deviation. *p<0.05, students t-test. C) Western blot analysis of phosphorylated Akt (threonine 342 and serine 505), phosphorylated S6K, and total actin (loading control), in lysates from mated adult flies after they had been exposed to either normoxia (N) or hypoxia (H) as larvae.

We next examined whether larval hypoxia exposure could alter adult TOR kinase signaling. One direct phosphorylation target of TOR is S6 kinase. Using an antibody that recognizes the TOR phosphorylation site on S6K, we found that, in contrast to insulin signaling, TOR activity was elevated in HYPOXIC adult flies (Figure 3C). Taken together, these data indicate that exposure of larvae to hypoxia leads to a long-term persistent suppression of insulin signalling but an increase in TOR signaling in adults.

### Larval hypoxia leads to sex-specific changes in adult nutrient storage

Animals often rely on mobilization of nutrient stores to fuel their metabolism during periods of stress, particularly nutrient deprivation. Studies in Drosophila have shown that genetic disruption of nutrient mobilization can reduce starvation tolerance and reduce lifespan (Mattila and Hietakangas, 2017; Heier and Kuhnlein, 2018). Since, we observed that larval hypoxia could exert effects on adult stress resistance and lifespan, we examined whether altered nutrient storage and mobilization might be involved. To do this, we analyzed adult whole body levels of triacylglycerides (TAG), the main lipid stores, and levels of glycogen, the main sugar stores, in the NORMOXIC and HYPOXIC conditions. These studies revealed a sexual dimorphic effect of larval hypoxia on adult nutrient stores. We found that the HYPOXIC condition males had significantly elevated levels of both TAG and glycogen compared to NORMOXIC males (Figure 4A). In contrast, we found that TAG and glycogen levels were unchanged between NORMOXIC and HYPOXIC females (Figure 4B). These findings that HYPOXIC animals have either normal (females) or elevated (males) levels of stored lipids and sugars are perhaps surprising given that these animals show reduced starvation tolerance compared to NORMOXIC animals. However, one possibility is that despite having high levels of stored nutrients, the HYPOXIC animals may have defects in nutrient mobilization. We therefore examined whether nutrient mobilization might be different between the NORMOXIC and HYPOXIC groups when they are subjected to nutrient starvation. We focused on looking at TAG since proper mobilization of lipid stores has been shown to be essential for starvation tolerance in Drosophila. Both NORMOXIC males and females showed a decrease in total TAG levels following starvation, with this effect being more pronounced in males, a previously reported result that is consistent with mobilization of lipid stores in nutrient deprived conditions (Gronke et al., 2007; Wat et al., 2020). In contrast, HYPOXIC animals showed a sexually dimorphic response to starvation. HYPOXIC males showed a significantly greater decrease in TAG levels upon starvation compared to NORMOXIC males, and in fact, almost complete depleted (94% decrease) their lipid stores (Figure 5). In contrast, HYPOXIC females did not show any decrease in TAG levels (Figure 5). Together, these data indicate that larval hypoxia exposure leads to abnormal nutrient storage and mobilization in adults.

**Fig 4.**
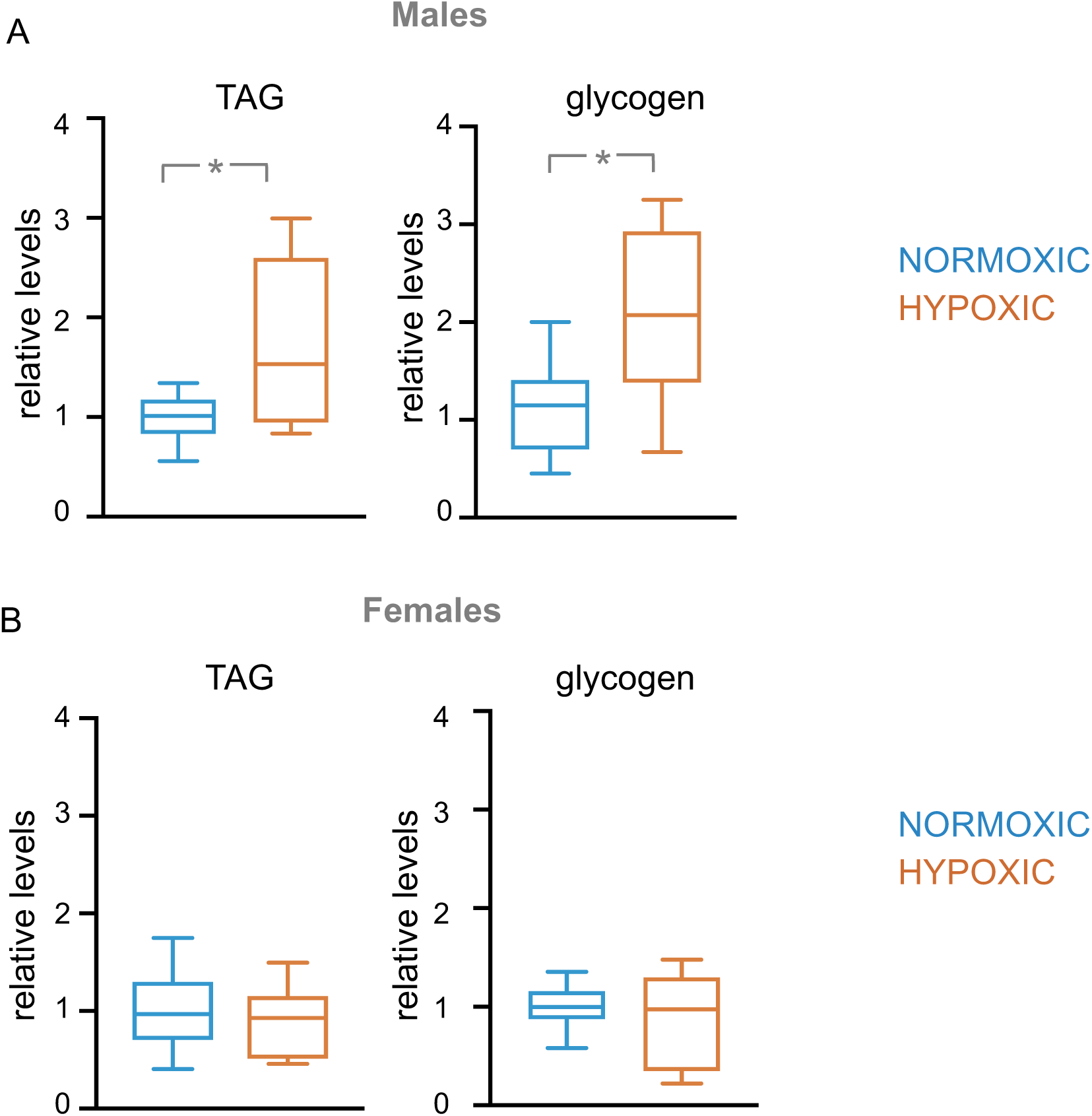
Larval hypoxia alters adult male, but not female, nutrient storage. Levels of TAG, glycogen, and glucose were measured in mated A) adult male and, B) female animals after they had been exposed to either normoxia (blue plots) or hypoxia (orange plots) as larvae. Data are presented as box plots (25%, median and 75% values) with error bars indicating the min and max values. N>10 per condition. *p<0.05, students t-test.

**Fig 5.**
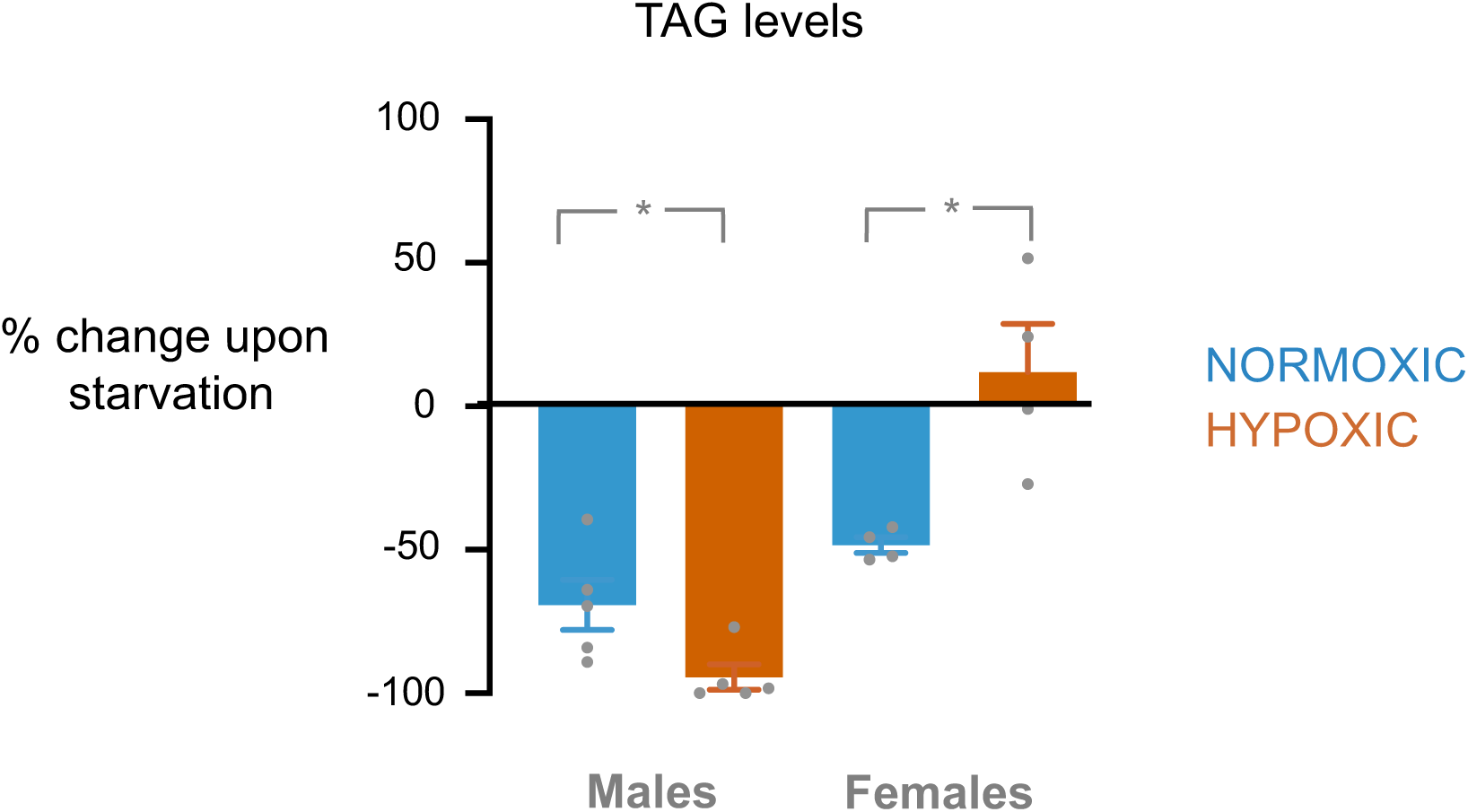
Larval hypoxia leads to a sexually dimorphic effect on lipid mobilization during starvation stress. Changes in TAG levels upon 16 hours of complete nutrient starvation in adult male and female animals after they had been exposed to either normoxia (blue bars) or hypoxia (orange bars) as larvae. Data are represented as mean percentage change in TAG levels +/-SEM. *p<0.05, students t-test.

## DISCUSSION

In this paper, we explored how early hypoxia affects adult physiology and homeostasis. In particular, we were interested in testing the possibility that early-life hypoxia might confer beneficial effects on adult fitness. However, we found that larval hypoxia exerted no hormetic effect of early life hypoxia on adults. This finding contrasts with previous studies that showed that two other larval environmental manipulations - nutrient restriction and oxidative stress – could extend adult lifespan (Stefana et al., 2017; Obata et al., 2018). Similarly, genetic manipulations that trigger a pulse of mitochondrial stress in the early larval period can also lead to extended lifespan in adults (Owusu-Ansah et al., 2013; Borch Jensen et al., 2017). A similar beneficial effect of early life mitochondrial stress has also been described in *C elegans* (Dillin et al., 2002). One possibility is that the type of early-life stress dictates later whether any beneficial effects are seen later in life. Thus, nutrient restriction and oxidative stress, but not oxygen limitation, may converge upon similar metabolic or regulatory processes to confer later effects on adult fitness. Alternatively, in the case of hypoxia, both the level and duration of larval low oxygen exposure may be important in determining whether any potential long-term effects in adults are beneficial or deleterious. A previous study showed that exposure of larvae to 10% oxygen also reduced adult lifespan (Rascon and Harrison, 2010). Thus it could be the case that exposure to less severe hypoxia, possibly just a little lower than the 20% oxygen level in air, could exert beneficial effects. Alternatively, a pulse of transient, more severe, but non-lethal, hypoxia or anoxia could be important. This has been seen in *C elegans* where short term 36h exposure to 0.5% oxygen leads to a subsequent extension of adult lifespan (Schieber and Chandel, 2014).

Rather than seeing beneficial effects we actually found that the effects of the larval hypoxia were deleterious to adult fitness – both male and female adults that had been exposed to hypoxia as larvae showed a reduced ability to tolerate nutrient starvation, and they had a shortened lifespan. Our metabolic analyses suggest that these effects could occur as a result of altered nutrient storage and mobilization caused by prior hypoxic exposure. Numerous studies have shown that the storage and mobilization of nutrient stores, particularly lipids, are essential for Drosophila to tolerate starvation stress and to maintain normal lifespan (Heier and Kuhnlein, 2018). This mobilization of nutrient stores is required to allow proper fuelling of key metabolic processes required to allow animals to survive in the absence of food (Gronke et al., 2007; Palanker et al., 2009; Molaei et al., 2019; Wat et al., 2020). In this context, we identified alterations in both lipid storage and mobilization in Drosophila adult previously exposed to hypoxia as larvae. Interestingly these alterations exhibited a sexual dimorphism. Firstly we saw that HYPOXIC males, but not females, had elevated levels of TAGs and glycogen. Given that nutrient stores are required for starvation survival, these high TAG and glycogen levels in HYPOXIC males might be expected to promote starvation survival rather than the reduction in starvation survival that we actually observed. However, we also found that both male and female HYPOXIC animals showed a defect in lipid mobilization upon starvation, and that these effects were sexually dimorphic. In the case of males, the NORMOXIC control group showed a reduction in total TAG levels following starvation - a phenotype that has been reported before and is consistent with mobilization of lipid stores to support survival during nutrient starvation (Gronke et al., 2007; Palanker et al., 2009; Wat et al., 2020). However, we found that this decreases in TAG levels was significantly exacerbated in the HYPOXIC group, who showed an almost complete depletion of their lipid stores. Thus the increased starvation death in this group could be because they deplete their lipids too rapidly. In contrast, we saw an opposite phenotype in females. While NORMOXIC females showed lipid mobilization following starvation, the HYPOXIC females showed no depletion in TAGs upon starvation. Thus, in contrast to HYPOXIC males, the HYPOXIC females probably show decreased starvation survival due to an inability to mobilize their lipid stores.

What mechanisms may explain this sexual dimorphic difference in metabolism? One possibility suggested by a recent study involves a sexually dimorphic regulation of the lipase, brummer. Wat et al showed that adult male flies showed higher starvation-mediated induction of brummer compared to females, and, as a result they showed faster lipid depletion and decreased starvation survival (Wat et al., 2020). In our case, it is possible that larval hypoxia may subsequently lead to male:female differences in brummer regulation that could explain why our HYPOXIC flies showed both sexual dimorphic changes in lipid mobilization and overall reduced starvation survival. Other studies have described how different transcriptional regulators and lipid metabolism genes coordinate lipid mobilization to promote starvation survival (Baker and Thummel, 2007; Heier and Kuhnlein, 2018). In some cases similar phenotypes to the ones we see are reported. For example, male flies mutant for the translational repressor, 4E-BP, show a similar phenotype to the male HYPOXIC flies in that they show an increased depletion of lipid stores and decreased survival upon starvation (Teleman et al., 2005). In contrast, flies mutant for the transcriptional regulator Sir4 display a phenotype similar to HYPOXIC females – they fail to mobilize lipid stores upon starvation and subsequently show reduced survival (Wood et al., 2018). Thus, it is possible that larval hypoxia may cause alterations in these regulatory genes to lead to adult metabolic phenotypes. However, most (almost all) studies looking at the mechanisms of lipid mobilization during nutrient starvation have described results in only one sex. Hence, it is unclear whether any of the underlying mechanisms reported in these studies exhibit sexual dimorphisms that might explain the male-female differences that we see.

Other possible mediators of the effects of larval hypoxia on adult metabolism and survival are alterations in insulin and TOR signaling. In *Drosophila*, these pathways are normally both induced by nutrient and oxygen availability and have been shown to be suppressed by starvation and hypoxia (Grewal, 2009; Wong et al., 2014; Lee et al., 2019; Texada et al., 2019). Interestingly, we found diverging effects on TOR and insulin signaling – whole body insulin signaling was lower while TOR activity was elevated. The effects on insulin signaling are probably explained by the decreased expression of dILP 2, 3 and 5, which are three main dILPS that are secreted from the brain and whose expression is altered by nutrient availability. The elevation in TOR activity might be due to altered expression of upstream signaling components. Do these alterations in insulin and TOR signaling explain why HYPOXIC animals have altered metabolic effects and reduced starvation survival and lifespan? Generally, lowering systemic insulin signaling confers extension of lifespan and tolerance to stress in Drosophila (Giannakou and Partridge, 2007). However, increased TOR signaling has been shown to reduce adult lifespan in *Drosophila* (Kapahi et al., 2004). Hence, in the case of early-life hypoxia exposure the increased TOR might dominate to control survival in the HYPOXIC animals. Moreover, although reduced insulin and increased TOR was seen in both males and female the flies, these changes could even potentially explain the dimorphism in lipid metabolic phenotypes that we observed. For example, a recent study showed that genetic suppression of insulin signaling in adult flies could alter sexually dimorphic differences in gene expression, including many metabolic genes (Graze et al., 2018). In addition, alterations in TOR signaling have also been shown to have sex-dependent effects in gene expression and on nutrient control of reproduction in *Drosophila* (Camus et al., 2019).

## ACKNOWLEDGEMENTS

Stocks obtained from the Bloomington *Drosophila* Stock Center (NIH P40OD018537) were used in this study. This work was supported by an NSERC Discovery grant to S.S.G. D.M.P was supported by an NSERC CGS-M graduate scholarship.

